# PERIOD phosphoclusters control temperature compensation of the *Drosophila* circadian clock

**DOI:** 10.1101/2021.12.23.474078

**Authors:** Radhika Joshi, Yao D. Cai, Yongliang Xia, Joanna C. Chiu, Patrick Emery

## Abstract

Temperature compensation is a critical feature of circadian rhythms, but how it is achieved remains elusive. Here, we uncovered the important role played by the *Drosophila* PERIOD (PER) phosphodegron in temperature compensation. Using CRISPR-Cas9, we introduced a series of mutations that altered three Serines (S44, 45 and 47) belonging to the PER phosphodegron, the functional homolog of mammalian PER2’s S487 phosphodegron, which impacts temperature compensation. While all three Serine to Alanine substitutions lengthened period at all temperatures tested, temperature compensation was differentially affected. S44A and S45A substitutions caused decreased temperature compensation, while S47A resulted in overcompensation. These results thus reveal unexpected functional heterogeneity of phosphodegron residues in thermal compensation. Furthermore, mutations impairing phosphorylation of the *per*^*s*^ phosphocluster decreased thermal compensation, consistent with its inhibitory role on S47 phosphorylation. Interestingly, the S47A substitution caused increased accumulation of hyper-phosphorylated PER at warmer temperatures. This finding was corroborated by cell culture assays in which S47A resulted in excessive temperature compensation of phosphorylation-dependent PER degradation. Thus, we show a novel role of the PER phosphodegron in temperature compensation through temperature-dependent modulation of the abundance of hyper-phosphorylated PER. Our work also reveals interesting mechanistic convergences and differences between mammalian and *Drosophila* temperature compensation of the circadian clock.

**Author summary:** Circadian rhythms are critical adaptive mechanisms that enable most organisms to adjust their physiology and behavior to the changes that occur in their environment every day. Ambient temperature varies constantly, but interestingly molecular circadian pacemakers do not accelerate with increasing temperature, while most biochemical reactions are sensitive to temperature. This phenomenon of circadian temperature compensation is poorly understood. Using genome editing and transgenic approaches, we found that two phosphorylated motifs in the Drosophila PERIOD protein, which regulate stability, impact temperature compensation. Moreover, we observed that mutation of a key Serine residue controlling PER degradation, S47, affects the accumulation of phosphorylated PER in a temperature-dependent manner, and causes PER degradation kinetics to become overly protected from increased temperature. As a result, the circadian clock of S47 mutant flies is excessively temperature-compensated. Our work thus reveals an interesting mechanism that controls temperature compensation in *Drosophila*. Moreover, comparison with mammals reveal interesting similarities, but also important differences in how temperature compensation of the circadian clock is achieved.

## Introduction

Circadian clocks are present across almost all life forms and allow them to adapt their activities with daily changes in their surrounding environment. In eukaryotes, the molecular circadian pacemaker is a self-sustained transcriptional feedback loop that repeats itself every ∼24 h, thus producing molecular, physiological and behavioral rhythms that closely match day length (1). In *Drosophila*, the circadian clock comprises four core components (2). The transcription factors CLOCK (CLK) and CYCLE (CYC) form a heterodimer that binds to E-box motifs upstream of the *period (per)* and *timeless (tim)* genes to promote their transcription. PER and TIM heterodimerize in the cytoplasm before entering the nucleus. Once inside the nucleus, the PER/TIM heterodimer binds to CLK/CYC and thus inhibits *per* and *tim* transcription. PER, TIM, and CLK undergo various post-translational modifications that affect the period of the clock, the most notable being the progressive phosphorylation driven by kinases such as DOUBLETIME (DBT), SHAGGY, CASEIN KINASE 1 alpha (CK1α) (3), CASEIN KINASE 2 (CK2) and NEMO (NMO) (4).

Circadian clocks exhibit three key properties (5). First, these clocks free-run with a period of ∼24h in the absence of external temporal cues (referred to as Zeitgebers) such as the daily light and temperature cycles. Second, these Zeitgebers can reset the clocks, thus anchoring the clock’s phase to the day/night cycle. Third, the period of circadian clocks is temperature-compensated. As ambient temperature increases, most enzymatic reactions speed up. However, under constant conditions, circadian clocks exhibit ∼24 h periodicities over a wide range of physiological temperatures, even in ectotherms. This was first observed by C. Pittendrigh while studying eclosion rhythms of *Drosophila pseudoobscura* (6).

The mechanism of temperature compensation is one of the long lasting mysteries of circadian rhythms. Different models have been proposed. For example, the network model proposes that multiple temperature-sensitive reactions cancel each other with changes in temperature. Early studies on Dinoflagellates indeed suggested that two temperature-sensitive reactions with opposite effects would help achieving constant period (7). At the biochemical level, phosphorylation mediated by CK1 and CK2 has received the most attention across species. Mutations in the β1 and α subunits of CK2 in *Neurospora* alter temperature compensation (8). Temperature compensation in *Arabidopsis* is modulated by the inhibitory effect of CK2 phosphorylation on CIRCADIAN ASSOCIATED CLOCK A1 (CCA1) binding. CCA1 binding to its target promoters increases with temperature, which is kept in check by the opposing effect of CK2 phosphorylation (9). In mammals, Isojima et al. (10) showed that the activity of CK1δ/ε enzyme on synthetic PER2 peptide is temperature-insensitive, thus providing buffering against temperature changes. Interestingly, the CK1ε^⍰au^ mutant shows an undercompensation phenotype (11,12). Furthermore, CK1δ/ε activity on PER2 results in a temperature-dependent phosphoswitch involving two phosphorylation sites (12–14). One of them is Serine (S)-478, also called phosphodegron. S478 phosphorylation causes βTrCP (beta-transducing repeat containing homolog protein)-dependent ubiquitination and degradation of mPER2 (13–15). The second is S659, the mouse homolog to the site of familial advanced sleep phase syndrome (FASPS) mutations (16). S659 phosphorylation inhibits phosphorylation of the phosphodegron, and thus PER2 degradation (12,17). S478 is preferentially phosphorylated at colder temperatures while FASPS domain phosphorylation increases at warmer temperatures, consequently balancing out PER stability (12,18). Along with phosphorylation, other processes such as sumoylation in plants (19), nuclear cytoplasmic ratio of PER (20) and TIM (21), PER intermolecular and PER-TIM interactions (22–24) have all been proposed to modulate temperature compensation of circadian rhythms.

Over the past several decades circadian mutants have fuelled our understanding of the molecular clockwork. Therefore, to gain better understanding into the mechanism of temperature compensation in *Drosophila* we turned towards *Drosophila* mutants known to have a temperature compensation defect. Interestingly, of the numerous mutations impacting circadian rhythm period, only a subset of mutants show a defect in temperature compensation (25–32). We were intrigued by the mutant *per*^*SLIH*^ (**S**ome **L**ike **I**t **H**ot, Figure 1A) due to its striking undercompensation phenotype of ∼1.5h when temperature is increased form 18°C to 29°C (31). The *per*^*SLIH*^ mutation causes a Serine (S) to Tyrosine (Y) substitution at the 45^th^ residue, which is a part of *Drosophila* PER’s phosphodegron (S44, 45 and 47) (33). The PER phosphodegron is regulated in a similar manner as its mammalian homolog (34)): it is phosphorylated by the CK1δ/ε homolog DBT (33,35,36), and its phosphorylation is inhibited by the *per*^*short*^ (*per*^*S*^) phosphocluster, which is equivalent to the mammalian FASPS domain (12). Phosphorylation events at the PER phosphodegron, S47 in particular, are critical for SLIMB-mediated PER degradation and thus period length (33,37,38). SLIMB is the fly homolog of βTrCP. Here, we show that, unexpectedly, the residues of the *Drosophila* phosphodegron are functionally heterogeneous and differentially modulate temperature compensation, even though they all lengthen circadian period. In addition, we found that the *per*^*s*^ phosphocluster, which negatively regulates S47 phosphorylation, also regulates temperature compensation. Furthermore, we show that substituting PER’s S47 residue to an Alanine (A) results in an overcompensated circadian period that correlate with increased hyper-phosphorylated PER levels in vivo at warm temperature and overcompensated PER degradation in a well-established cell culture model. We therefore propose that PER phosphorylation and subsequent degradation is critical to temperature compensation in *Drosophila*.

**Figure 1:**
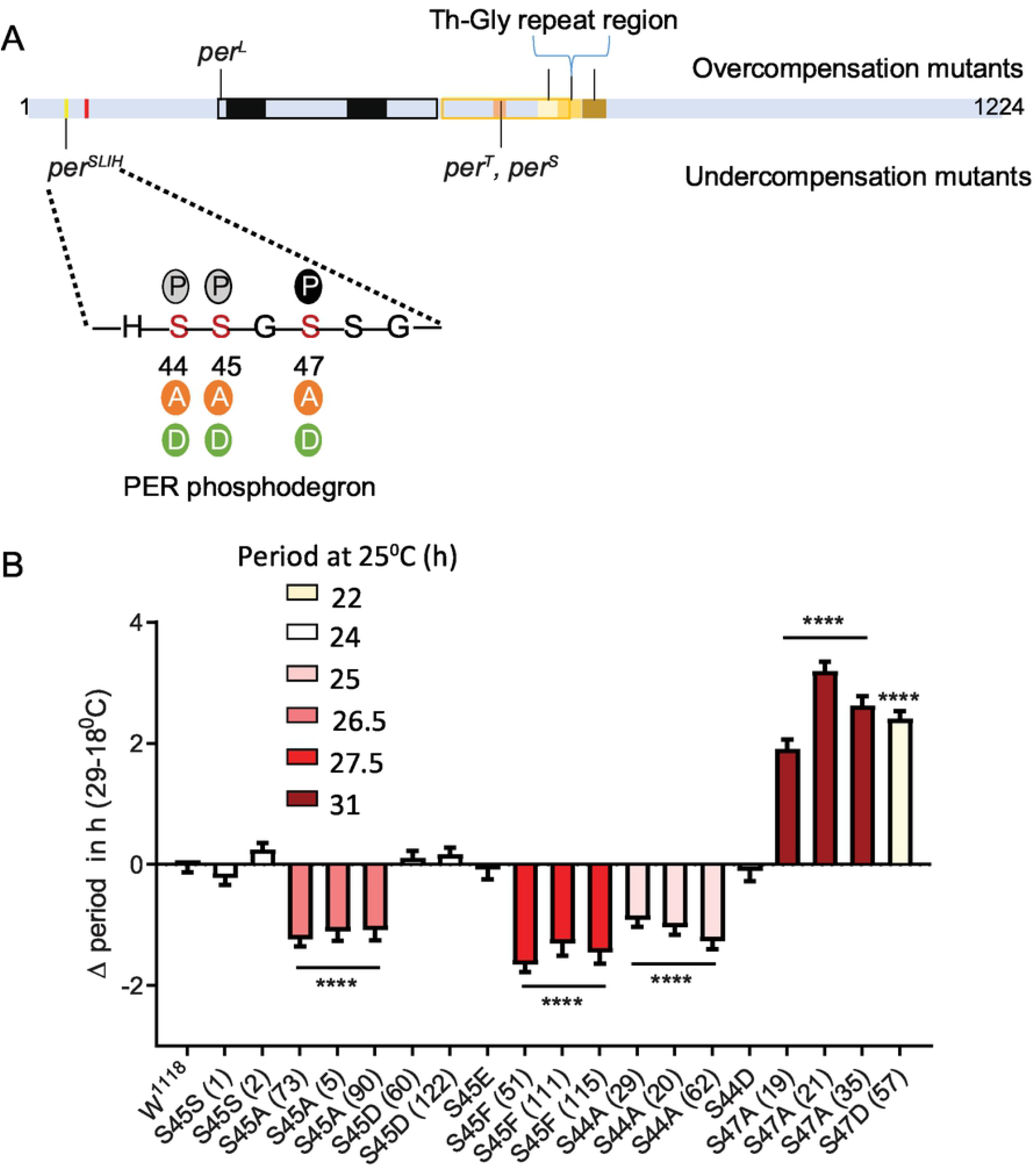
Differential modulation of temperature compensation by three Serine residues in the PER phosphodegron. A) Schematic of PER protein depicting the relative positions of amino acids modified in various mutants affecting temperature compensation in *Drosophila*. Mutants above the PER schematic are overcompensated while mutants depicted below are undercompensated mutants. The magnified region of PER shows residues of the dPER phosphodegron and the various substitutions (Serine to Alanine, Aspartate or Glutamate) we generated using CRISPR/CAS9. Solid P within black circle, shows that S47 position undergoes phosphorylation. Grey circles indicate potential additional phosphorylation sites at S44 and S45 (33). B) Graphical representation of the difference in period values of various mutants of dPER phosphodegron at 29^0^C and 18^0^C, relative to controls. ΔPeriod values for each genotype were offset by the average difference in period observed at 29^0^C and 18^0^C in S45SI and S45SII control lines, so that overcompensation or undercompensation compared to control is clearly represented. The color scheme denotes period of each mutant line at 25^0^C. Interestingly, S44A, S45A and S47A all produce long period phenotype at 25^0^C. However, the directionality of their temperature compensation phenotypes is opposite between S44A/S45A and S47A. S44A and S45A produce significant undercompensation while S47A and S47D shows overcompensation. Statistical significance was calculated using two-way ANOVA with Sidak’s multiple comparison test.

## Results

### Differential impact of three PER phosphodegron residues on temperature compensation

The unique location of the *per*^*SLIH*^ mutation (Figure 1A) prompted us to evaluate the role of residues of the PER phosphodegron in temperature compensation. DBT, a Serine/Threonine kinase, phosphorylates S47, with S44 and S45 being additional potential targets (33). Thus, it would seem likely that phosphorylation at these residues modulate temperature compensation. To explore this possibility, we made phosphomimics (S to D (Aspartate) or E (Glutamate)) and phosphoinhibitors (S to Alanine (A)) at the PER S44, S45, and S47 using CRISPR-CAS9 (Figure 1A). Moreover, we generated S45F (S to Phenylalanine) to mimic contribution of a bulky hydrophobic amino acid similar to Tyrosine (Y) in *per*^*SLIH*^. We used S45S as a control, which contains only the silent substitutions introduced at the protospacer adjacent motifs (PAM site) during CRISPR mutagenesis. In our hands *w*^*1118*^ and S45S control strains showed a decrease in period of ∼0.9h (Table 1) over the range of 18 to 29^0^C. To accurately represent the effect of mutations on temperature compensation, values plotted on figure 1B are relative to the average of the two S45S control strains, which was set to 0.

**Table 1:** Circadian behavior of control and *per* mutant flies under different temperatures. Mutations in the phosphoclusters were generated by CRISPR/Cas9-mediated genome editing. Amino acid substituions are indicated in the left column. *w*^*1118*^ is a wild-type strain. For the *per*^*S*^ phosphocluster, *per*^*0*^ flies were rescued with various transgenes. 13.2WT carries a wild-type per genomic region, while the other transgenes carry the indicated subsitutions. AR = arrhythmic ND = not done

| Degree Celsius | Period (h) |  |  |  | % Rhythmicity |  |  |  | # Rhythmic flies |  |  |  |
| --- | --- | --- | --- | --- | --- | --- | --- | --- | --- | --- | --- | --- |
|  | 29 | 25 | 20 | 18 | 29 | 25 | 20 | 18 | 29 | 25 | 20 | 18 |
| **Phosphodegrogen cluster** |  |  |  |  |  |  |  |  |  |  |  |  |
| W1118 | 23.6 | 24.5 | 24.3 | 24.5 | 87.5 | 86.9 | 53.8 | 40.0 | 93 | 53 | 42 | 32 |
| S45S (1) | 23.8 | 25.0 | 24.8 | 25.1 | 80.7 | 95.7 | 76.6 | 76.6 | 71 | 44 | 85 | 36 |
| S45S (2) | 23.9 | 24.9 | 24.8 | 24.7 | 83.3 | 95.3 | 61.0 | 84.7 | 60 | 61 | 64 | 50 |
| S45F (51) | 26.6 | 27.7 | 28.7 | 29.2 | 62.9 | 83.9 | 71.3 | 87.1 | 34 | 26 | 67 | 54 |
| S45F(111) | 26.4 | 27.3 | 28.2 | 28.7 | 60.0 | 53.3 | 65.8 | 37.5 | 18 | 8 | 25 | 12 |
| S45F(115) | 26.2 | 27.5 | 28.3 | 28.7 | 62.9 | 90.9 | 77.8 | 67.7 | 17 | 20 | 35 | 21 |
| S45A (73) | 26.3 | 27.2 | 28.0 | 28.5 | 76.6 | 93.5 | 82.1 | 87.5 | 36 | 29 | 78 | 56 |
| S45A (5) | 26.0 | 26.7 | 27.2 | 28.1 | 80.4 | 62.5 | 36.6 | 60.0 | 37 | 10 | 34 | 18 |
| S45A (90) | 26.3 | 26.8 | 27.8 | 28.4 | 65.0 | 93.8 | 75.4 | 70.0 | 26 | 15 | 46 | 21 |
| S45D (67) | 24.1 | 24.9 | 24.8 | 24.9 | 84.4 | 100.0 | 76.0 | 95.1 | 38 | 32 | 73 | 58 |
| S45D (122) | 24.0 | 24.9 | 24.8 | 24.8 | 84.1 | 95.1 | 80.0 | 91.1 | 53 | 58 | 88 | 51 |
| S45E | 23.9 | 24.7 | 24.2 | 24.9 | 87.2 | 93.3 | 44.2 | 35.4 | 41 | 43 | 19 | 17 |
| S44A (29) | 24.9 | 25.8 | 25.7 | 26.8 | 88.6 | 97.5 | 82.1 | 73.9 | 70 | 39 | 69 | 34 |
| S44A (20) | 24.9 | 26.0 | 26.6 | 26.9 | 77.1 | 91.3 | 84.8 | 89.1 | 37 | 21 | 56 | 41 |
| S44A (62) | 24.7 | 24.2 | ND | 27.0 | 76.7 | 75.0 | ND | 55.0 | 33 | 14 | ND | 33 |
| S44D | 25.6 | 26.6 | 26.9 | 26.7 | 62.5 | 90.7 | 66.7 | 35.7 | 25 | 39 | 29 | 20 |
| S47A (19) | 32.1 | 31.3 | 31.0 | 31.2 | 28.8 | 53.8 | 46.9 | 52.1 | 17 | 44 | 15 | 49 |
| S47A (21) | 33.5 | 31.5 | 31.5 | 31.3 | 38.3 | 44.9 | 15.6 | 58.8 | 18 | 48 | 5 | 47 |
| S47A (35) | 33.0 | 31.5 | 31.3 | 31.4 | 23.3 | 72.9 | 37.5 | 47.7 | 17 | 70 | 12 | 53 |
| S47D | 22.5 | 22.6 | 21.7 | 21.1 | 88.5 | 97.9 | 93.6 | 72.7 | 46 | 47 | 44 | 40 |
| S4445A (105) | 34.4 | 35.9 | 35.3 | 38.0 | 56.3 | 32.3 | 35.6 | 6.3 | 9 | 10 | 16 | 1 |
| S4445A (32) | 34.0 | AR | 35.6 | AR | 19.4 | 0.0 | 4.2 | 0.0 | 6 | 0 | 2 | 0 |
| S4445A (60) | 34.1 | AR | 32.2 | AR | 59.4 | 0.0 | 25.4 | 0.0 | 19 | 0 | 16 | 0 |
| S4547A (1) | AR | AR | AR | AR | 0.0 | 0.0 | 0.0 | 0.0 |  |  |  |  |
| S4547A (30) | AR | AR | AR | AR | 0.0 | 0.0 | 0.0 | 0.0 |  |  |  |  |
| S4547A (57) | AR | AR | AR | AR | 0.0 | 0.0 | 0.0 | 0.0 |  |  |  |  |
| S4547A (84) | AR | AR | AR | AR | 0.0 | 0.0 | 0.0 | 0.0 |  |  |  |  |
| S4547A (90) | AR | AR | AR | AR | 0.0 | 0.0 | 0.0 | 0.0 |  |  |  |  |
| **per<sup>s</sup> cluster** |  |  |  |  |  |  |  |  |  |  |  |  |
| 13.2WT | 23.8 | 24.2 | 23.9 | 23.8 | 89.3 | 76.6 | 23.3 | 32.9 | 75 | 37 | 15 | 27 |
| S47A (M28-F1) | 32.7 | 30.5 | 30.0 | 29.9 | 48.3 | 44.7 | 9.9 | 27.2 | 43 | 21 | 9 | 28 |
| S47A (F15-F1) | 32.6 | 31.6 | 31.4 | 29.8 | 96.8 | 95.8 | 65.6 | 50.8 | 61 | 46 | 43 | 32 |
| S47D (M7-M2) | 21.7 | 21.8 | 21.5 | 21.0 | 69.6 | 80.4 | 68.1 | 44.8 | 48 | 37 | 47 | 23 |
| S596A (F1-F1) | 15.9 | 15.2 | 15.9 | 16.3 | 75.0 | 91.5 | 18.8 | 8.5 | 33 | 43 | 9 | 4 |
| S596A (M16-F1) | 15.7 | 15.5 | 16.8 | 17.2 | 68.1 | 93.8 | 95.7 | 81.6 | 32 | 45 | 44 | 62 |
| TS583-596A (F30-M1) | 16.9 | 17.4 | 17.7 | 17.8 | 22.9 | 72.3 | 60.4 | 67.3 | 11 | 34 | 29 | 37 |
| TS583-596A (M3-F1) | 16.2 | 16.3 | 17.0 | 17.3 | 61.7 | 80.9 | 82.6 | 86.8 | 29 | 38 | 24 | 79 |
| TS583-596D (F2-F1) | 15.3 | 15.0 | 15.8 | 16.0 | 55.3 | 91.5 | 45.8 | 48.6 | 26 | 43 | 22 | 41 |
| TS583-596D (F2-M1) | 15.1 | 15.2 | 15.7 | 16.0 | 68.8 | 78.6 | 69.0 | 60.0 | 22 | 22 | 20 | 37 |
| S589A (M8-F1) | 19.4 | 19.6 | 21.0 | 20.7 | 95.1 | 97.9 | 97.9 | 76.7 | 39 | 46 | 46 | 56 |

Consistent with previous observations (33), we observed that the phosphoinhibitory substitution S47A produced lengthened period. In addition, we found that S44A, S45A, S45F also resulted in longer periods, albeit to different extent at 25^0^C (Figure 1B, Table 1). S44A lengthens the period by ∼1.5h, S45A by 2.5h, and S47A by 6.5h at 25^0^C. Interestingly, the effect on temperature compensation did not correlate with period length. S44A and S45A/F were undercompensated by ∼1 to 2h, compared to controls. However, S47A showed an opposite phenotype: it caused a strong overcompensation of more than 2h (Figure 1B). Since it is possible that mutations at a residue affect phosphorylation events at the neighboring residues, we generated the double mutants S44-45A and S45-47A. S44-45A double mutants exhibited a very long period (>34h) at 25^0^C. Period was very long, but the effect on temperature compensation was difficult to interpret due to poor rhythmicity, particularly at 18^0^C (Table 1). However, the synergistic effects on period and rhythmicity indicate that S44 and 45 work together to control circadian rhythms. S45-47A mutants were completely arrhythmic at both temperatures tested (Table 1).

We also tested phosphomimic substitutions (S to D/E). S45D and S45E did not show any substantial effect on either period length or temperature compensation (Figure 1, Table 1). S44D lengthened the period to 26.6h at 25^0^C, but had no effect on temperature compensation The behavior of S47D was the most peculiar. It showed a short rhythm period phenotype (>21h), as previously reported by Chiu et al (33) with *per* genomic rescue constructs in a *per*^*0*^ background. However, period was overcompensated by more than 2h. This could either be because phosphomimics do not correctly reproduce the structure of a phosphorylated residue (only its charge), or because phosphomimics are neither temporally controlled nor reversible and could thus have unexpected impact on temperature compensation.

Thus, taken together our results show that the PER phosphodegron modulates temperature compensation, and shows functional temperature compensation heterogeneity within its residues.

### Role of the *per*^*s*^ domain in temperature compensation

To further strengthen evidence for a role of phosphorylation at the PER phosphodegron in temperature compensation, we analyzed the impact of mutations in the *per*^*s*^ domain (T/S583-596). Indeed, this region also undergoes phosphorylation by DBT and NMO, and phosphorylation in the *per*^*s*^ domain temporally precedes and inhibits phosphorylation at S47 (39). Therefore we tested point mutations in the *per*^*s*^ domain for their role in temperature compensation.

We used previously described transgenic lines (39), in which the *per*^*0*^ mutation is rescued with *per* transgenes carrying various mutations in the *per*^*s*^ domain. In addition, we included transgenic flies carrying S47A and S47D substitutions. Strikingly, the period length and overcompensation phenotypes observed with transgenes encoding PER S47A and S47D were similar to those observed with the corresponding mutants generated with CRISPR-CAS9 (Figure 1 and 2), confirming the critical role played by the S47 residue in temperature compensation. Mutants in the *per*^*s*^ domain exhibited shortened period length at 25^0^C (as described previously (39). They also showed consistently an undercompensation phenotype of about 1-1.5 hours, except for one S596A line (Figure 2B). This could be because of an insertional effect, since *per* transgenes were randomly inserted in the fly genome. It should be noted that the *per*^*T*^ and *per*^*S*^ mutations have also been reported to be undercompensated (28–30). Taken together, our results and these previous reports show that the *per*^*S*^ domain is important for temperature compensation. Since S to A substitutions in this domain have opposite effects than S47A, and since phosphorylation of the *per*^*S*^ domain blocks S47 phosphorylation, it seems likely that the *per*^*S*^ domain controls temperature compensation through S47 phosphorylation.

**Figure 2:**
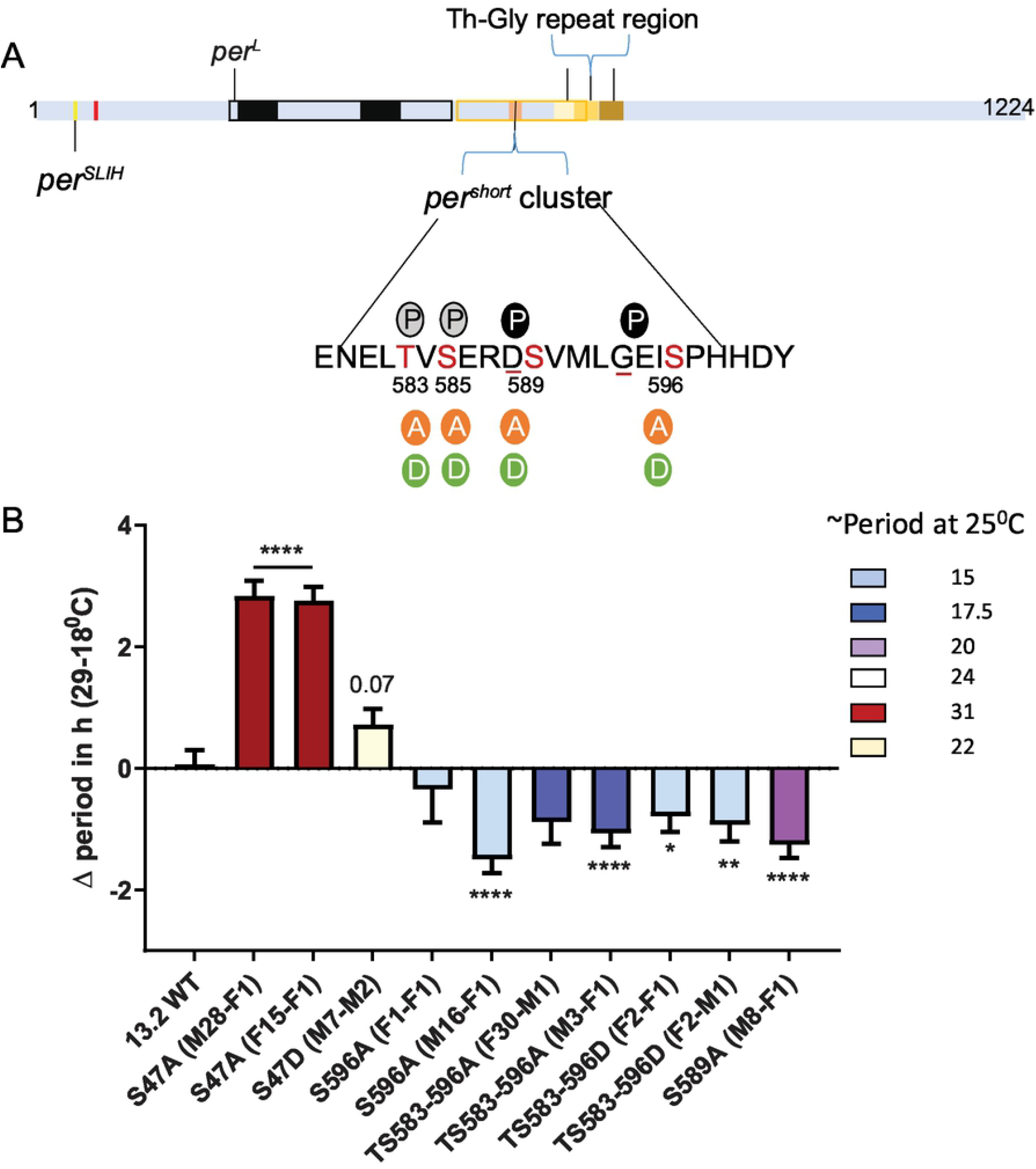
the *per*^*S*^ phosphocluster impacts temperature compensation. A) PER schematic depicting the relative position of various mutants affecting temperature compensation, as shown in figure 1A. The magnified region shows key Serine residues of the *per*^*S*^ phosphocluster and the various substitutions that were tested. Solid P within black circle at 589 and 596 shows sites of DBT and NMO mediated phosphorylation, respectively. Dotted grey circles show predicted phosphorylation sites at S585 and S583. Underlines at S589 and G593 denote positions of *per*^*S*^ and *per*^*T*^ mutant, respectively. B) Graphical representation of the difference in period of various mutants of the *per*^*s*^ domain at 29^0^C and 18^0^C. The color scheme denotes period of each mutant line at 25^0^C. All mutants in the *per*^*s*^ domain produce short period phenotype at 25^0^C. Most show significant undercompensation. Transgenic lines with S47A recapitulated our observations with CRISPR-Cas9 genome editing (figure 1). Transgenic S47D showed a weaker phenotype than genome edited S47D, just short of statistical significance (P=0.07). Statistical significance was calculated using two-way ANOVA with Sidak’s multiple comparison test.

### S47A mutants show increased hyper-phosphorylated PER accumulation at warmer temperatures

To further understand the mechanism of temperature compensation, we examined PER protein cycling in control, S45A, and S47A mutants at 18^0^C and 29^0^C in head extracts. Under LD, control (*w*^*1118*^) flies showed rhythmic PER abundance and phosphorylation regardless of the temperature, albeit with significantly higher abundance at warmer temperatures (Figure 3A, B and C). This observation is in line with increased TIM abundance at warm temperature (40).

**Figure 3:**
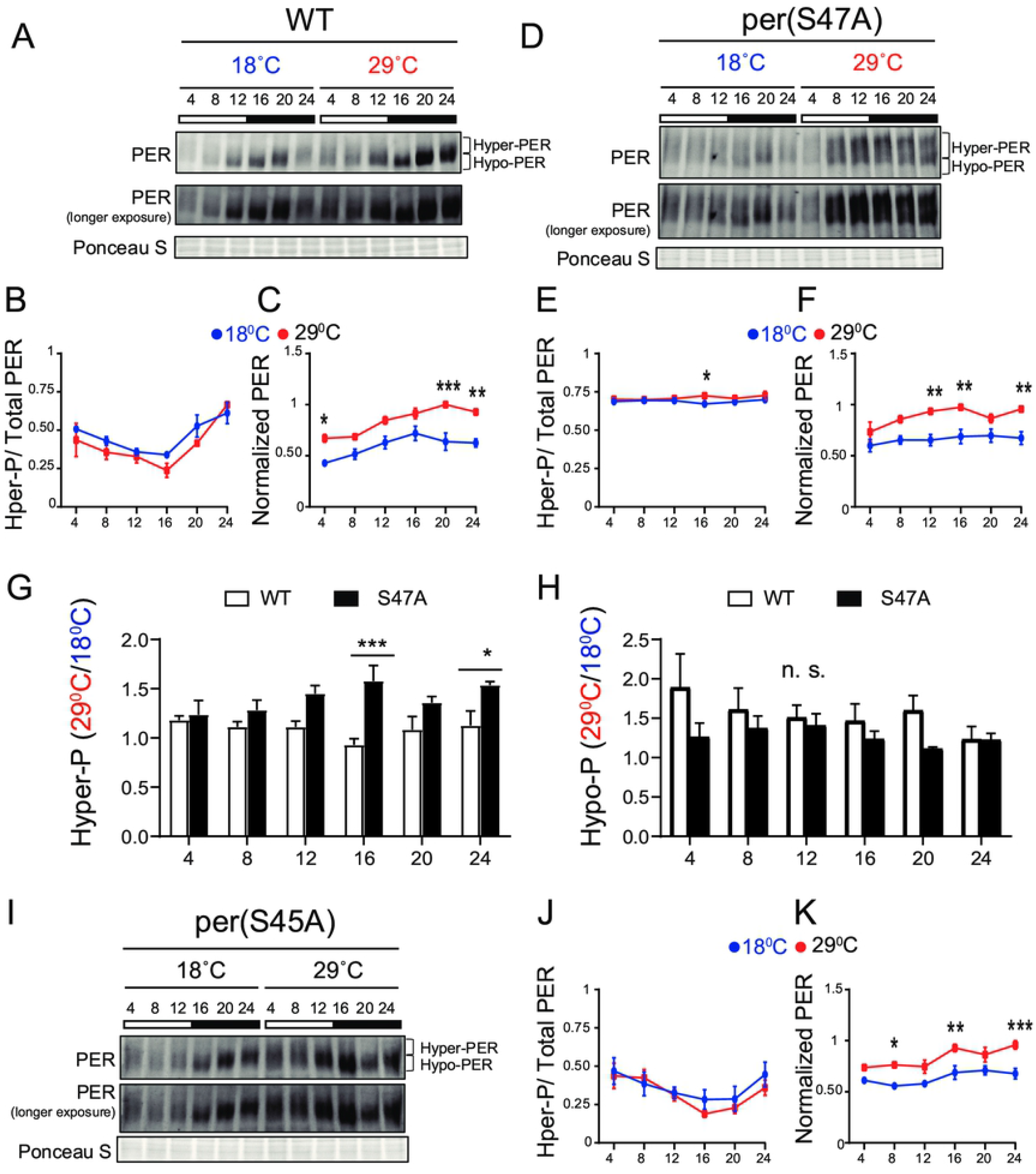
S47A mutation causes hyper-phosphorylated PER to accumulate preferentially at high temperature. A, D and I) Representative western blots probing head extracts from flies entrained to LD at 18^0^C and 29^0^C. PER abundance was monitored throughout the day at the indicated time points. Respective genotypes are represented at the top of each gel. Brackets indicate hyper-and hypo-phosphorylated PER isoforms. Two different exposures of the same blot are shown. Ponceau S was used as a loading control and for normalization. B-F and J-K) Graphs show PER abundance and phosphorylation quantifications. In control flies, both PER phosphorylation (B) and abundance (C) were scored as statistically rhythmic at 29^0^C and 18^0^C, using the JTK_CYCLE. For phosphorylation: p<0.05, phase=2 at 18^0^C and p<0.001, phase=2 at 29^0^C. For abundance: p<0.05, phase=18 at 18^0^C and p<0.001, phase=20 at 29^0^C. For S47A, neither phosphorylation (E) nor abundance (F) were scored as statistically rhythmic, although PER levels were reproducibly lower at ZT4. For S45A, only PER abundance at 18^0^C scored as statistically rhythmic. p<0.05, phase=18. Red lines represent data at 29^0^C and blue lines represent data at 18^0^C. *p<0.05, **p<0.01, ***p<0.001, two-way ANOVA followed by Sidak’s multiple comparison to test for time and temperature-dependent differences (n=3 biological replicates). G-H) Ratio of PER levels measured at 29^0^C vs 18^0^C. G shows the ratio for hyper-phosphorylated PER, and H for hypo-phosphorylated PER. S47A accumulates more hyper-phosphorylated PER during the night than control flies at warm temperatures.

S47A also showed increased PER abundance at 29^0^C compared to 18^0^C, but PER abundance and phosphorylation rhythms were perturbed at both temperature (Figure 3D, E F). Importantly, S47A mutants showed exaggerated temperature-sensitive accumulation of hyper-phosphorylated PER compared to controls at night (Figure 3G), but not of hypo-phosphorylated PER (Figure 3H). Increased accumulation of hyper-phosphorylated PER is likely to result in longer repression phase thus lengthening the period length (41) at warmer temperatures. These results therefore reveal a temperature-dependent effect on PER phosphorylation and/or degradation of the S47A substitution, which could underlie the observed overcompensation phenotype.

S45A showed a milder phenotype than S47A, with patterns of phosphorylation similar to that of WT (Figure 3 I, J and K). However, PER cycling amplitude at warmer temperatures was reduced and did not clear statistical significance. Phosphorylation rhythm amplitude were reduced in amplitude at both temperature as well.

### PER degradation is overcompensated as a result of the S47A substitution

The increase in the accumulation of hyper-phosphorylated PER, observed at high temperature with the S47A substitution, could be the result of abnormal protein stability, possibly due to abnormal phosphorylation. We therefore turned to validated S2 cell culture assays (33) to determine the effect of temperature on PER phosphorylation and degradation kinetics. In these assays, recombinant DBT is expressed under the inducible metallothionine promoter, while PER is expressed constitutively. Since PER is phosphorylated on multiple sites by DBT and thus produce several phosphorylated isoforms with various mobility that are subjected to SLIMB-mediated proteasomal degradation, we quantified the disappearance of the hypo-phosphorylated PER after DBT induction to determine PER phosphorylation kinetics (Figure 4). Interestingly, phosphorylation kinetics of PER S47A was significantly slowed down 6 hours following DBT induction at low temperature (Figure 4A, C), while no significant effect was observed with either wild-type PER or S45A at this time point (Figure 4A, B and D). However, it seems unlikely that a slower phosphorylation kinetic at cold temperature would result in the shorter circadian behavior period observed in S47A mutants at 18°C compared to 29°C.

**Figure 4:**
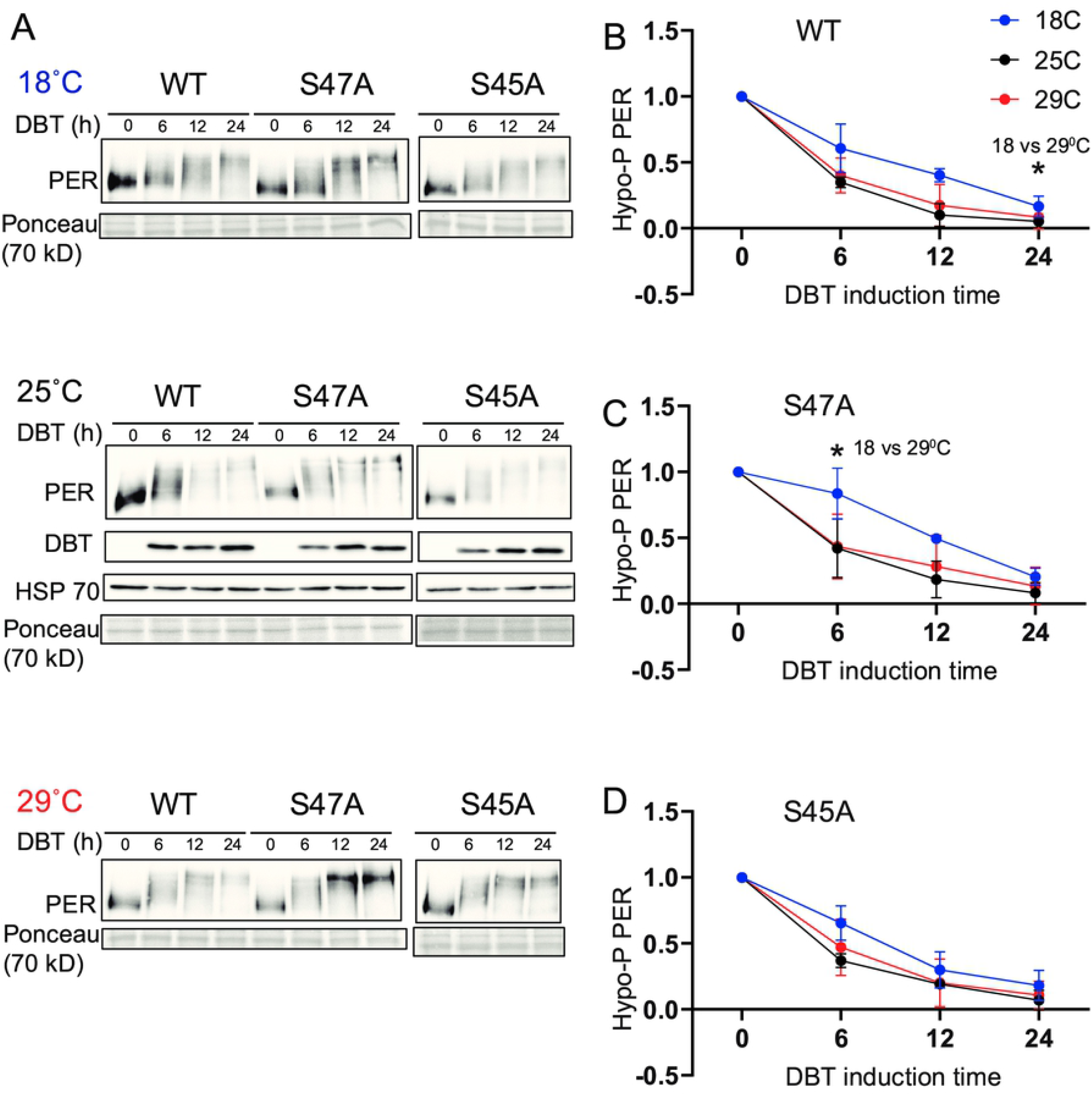
S47A phosphorylation kinetics is slowed down at cold temperature. Representative western blots probing cell extracts from S2 cells expressing DBT and either wild-type (WT) PER, S47A or S45A. Cells were incubated at the indicated temperatures, and collected at the indicated time points after DBT induction. Top panel shows PER immunoblotting, bottom panel shows Ponceau S staining, used as loading control and for normalization. B, C and D) Western Blot quantifications. PER signal shown at time point 0 is used as a reference to classify PER isoforms as hypo-phosphorylated. Graphs representing the relative amount of hypo-phosphorylated PER at 18, 25 and 29^0^C for the respective genotypes at different time points after DBT induction. S47A shows increased hypo-phosphorylated PER 6h after DBT induction.*p<0.05, two-way ANOVA followed by Sidak’s multiple comparison to test for time and temperature-dependent differences (n=3 biological replicates).

Since S47A caused increased hyper-phosphorylated PER accumulation at 29^0^C, we also determined S47A’s effects on PER stability. We again induced DBT and after 6 hours added cycloheximide to block protein synthesis. Wild-type PER was much more stable than S47A at 18^0^C, while at 29^0^C there was no obvious difference (Figure 5B and C). Strikingly, S47A’s degradation kinetics was slowed down at higher temperature, while for WT, it was accelerated (Figure 5A, D, E). Thus, S47A results in a thermal overcompensation of PER degradation, which probably contribute to the excessive hyper-phosphorylated PER visible in flies expressing S47A, and thus causes overcompensation of circadian period (Figure 6).

**Figure 5:**
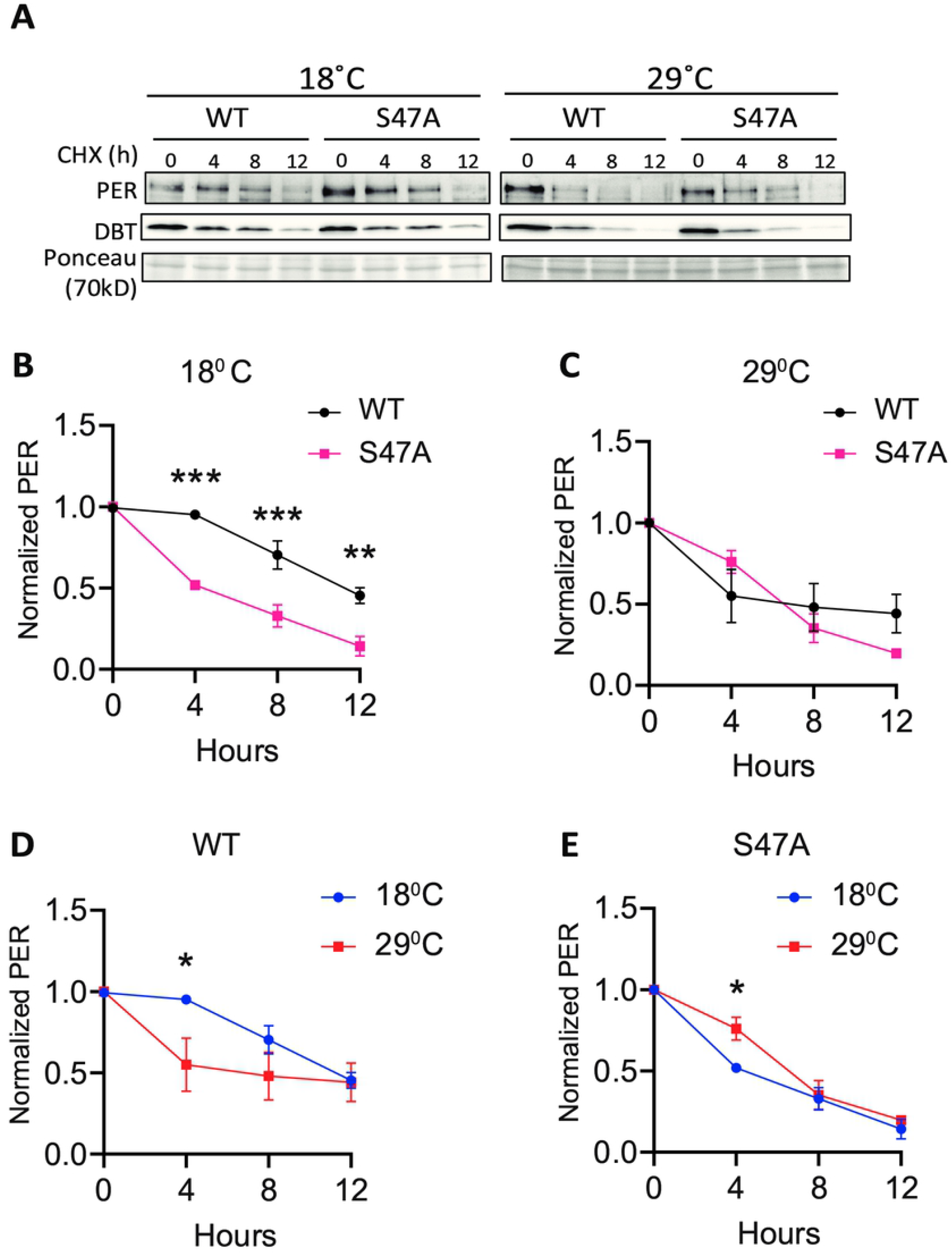
S47A degradation kinetics is excessively temperature-compensated. A) Representative western blots probing cell extracts from S2 cells expressing DBT and either wild-type (WT) PER, S47A or S45A. Cells were collected at the indicated time points after cycloheximide addition at the indicated temperatures. Top panel shows PER immunoblotting, bottom panel shows Ponceau S staining, used as loading control and for normalization. B-E) Western Blot quantifications. All experiments were performed with three independent replicates. *p<0.05, **p<0.01, ***p<0.001, two-way ANOVA followed by Sidak’s multiple comparison to test for time and temperature-dependent differences. In B and C, WT and S47A mutants are compared to each other at the indicated temperature. D and E show the same data, but degradation kinetics of WT and S47A are compared between high and low temperatures.

**Figure 6:**
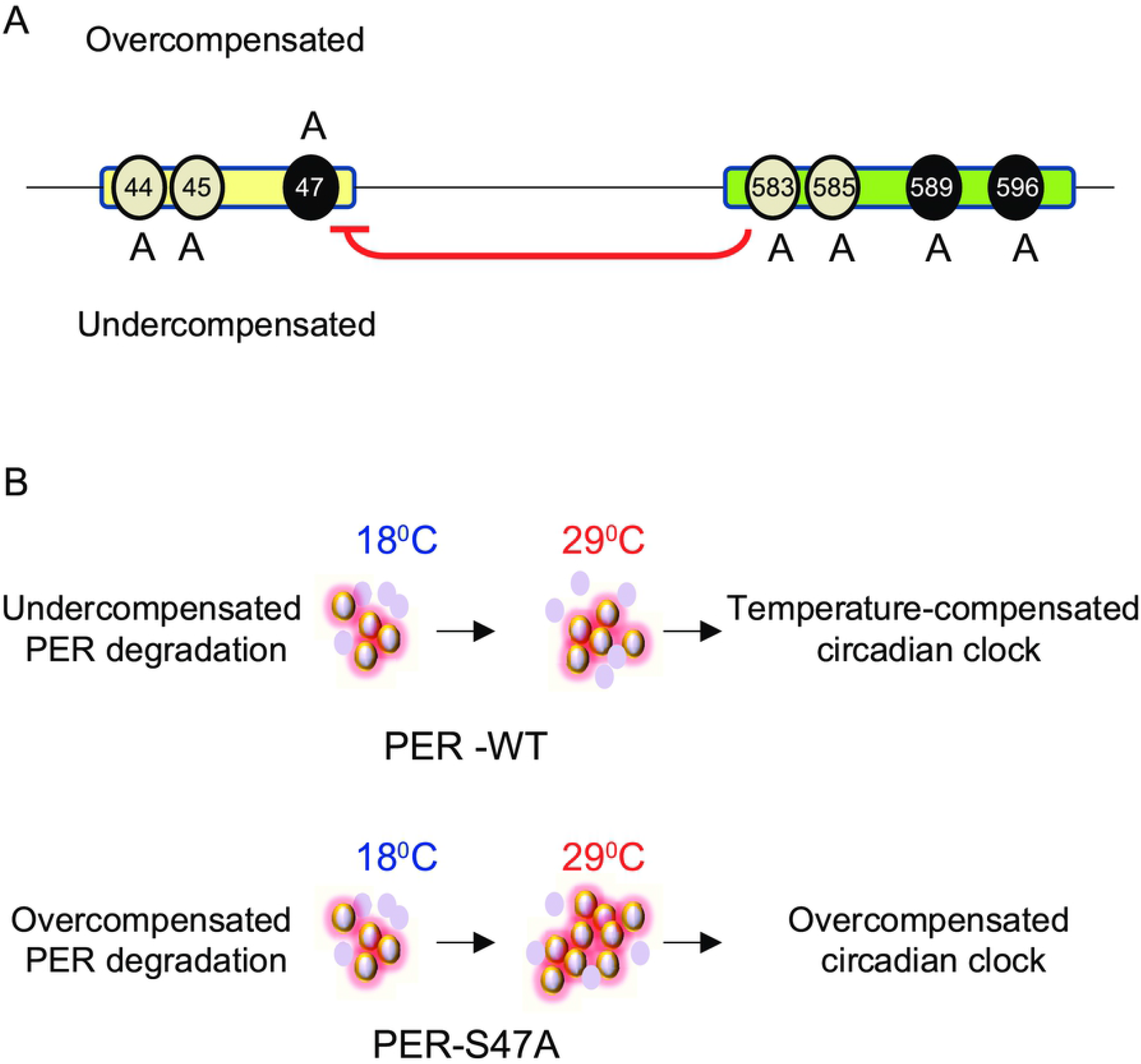
Model for the role of PER phosphoclusters in temperature compensation. A) The PER phosphodegron (yellow) and *per*^*s*^ phosphocluster (green) modulate temperature compensation. Mutations of different residues of the PER phosphodegron have opposite effects on temperature compensation. S44A and S45A cause undercompensation while S47A results in overcompensation of circadian period. Mutations in the *per*^*s*^ phosphocluster cause undercompensation, consistent with its inhibitory role on S47 phosphorylation (red arrow). Black ovals around 47, 589, and 596 show DBT and NMO mediated phosphorylation sites on PER. Grey ovals around 44, 45, 583, and 585 depict putative additional phosphorylation sites. B) Mechanism of S47 action on temperature compensation. In wild-type flies, S47 phosphorylation causes PER degradation to speed up as temperature increases. Hyper- and hypo-phosphorylated isoforms are in balance and the clock is properly temperature-compensated. In S47A mutant flies, PER degradation is overcompensated and thus slows down at high temperature, leading to excessive accumulation of hyperphosphorylated isoforms. The clock is then overcompensated. Violet ovals represent PER protein, Halo around violet ovals show hyper-phosphorylated PER.

## Discussion

Phosphorylation events play an important role in the temperature compensation of the circadian clock across phyla (8,9,11–13). In *Drosophila*, PER phosphorylation has been studied in exquisite detail, and interestingly three *per* mutations affecting two PER domains undergoing DBT phosphorylation are undercompensated. *per*^*SLIH*^ causes the most robust undercompensation phenotype, while *per*^*S*^ and *per*^*T*^ have milder effects (25,28–30). We therefore systematically tested phosphorylated residues in both the *per*^*S*^ domain and the PER phosphodegron. Overall, our results point at a central role for residue S47, and modulatory roles for S44, S45 and residue in the *per*^*S*^ phosphorylation cluster.

Indeed, the S47A substitution resulted in a striking increase in temperature compensation. This suggests that phosphorylation at this residue is important for the compensation. Actually, we found that PER levels were exaggeratedly temperature-dependent *in vivo*, with a striking increase in hyper-phosphorylated PER levels at high temperature. Moreover, PER degradation induced by DBT phosphorylation was slowed down at high temperature, rather than accelerated by the S47A substitution in a *Drosophila* cell culture assay. These results thus suggest that S47 phosphorylation controls the kinetics of PER degradation in a temperature-dependent manner, protecting degradation from being overcompensated. This is further supported by the observation that mutations in the *per*^*S*^ phosphocluster, which inhibits S47 phosphorylation, have opposite effects on temperature compensation: flies carrying these mutations are undercompensated. Unexpectedly though, the phosphomimetic substitution S47D did not produce an undercompensation phenotype. Rather, an overcompensation phenotype was observed. This could be explained by the substitution to Aspartate not properly mimicking the structure of a phosphorylated Serine, or that constantly mimicking S47 phosphorylation impacts thermal compensation of the clock through a different mechanism than the S47A substitution.

Another unexpected observation is that Serine to Alanine substitution at residues 44 and 45 have opposite effects on temperature compensation than S47A, even though all these substitutions lengthen circadian period. Residues of the phosphodegron are thus functionally heterogeneous when it comes to temperature compensation. Indeed, unlike S47A, S45A did not cause obvious change in temperature-dependent PER stability *in vivo* or in cell culture. It should be noted that unlike S47, S44 or S45 phosphorylation by DBT has not been unambiguously demonstrated (33). It remains thus possible that these residues are not phosphorylated and play a structural role in the phosphodegron, and this could explain functional diversity in temperature compensation between S44/45 and S47. Further studies are needed to elucidate S44 and S45’s exact function in thermal compensation. Of note, the S45F substitution, which introduces a more bulky amino acid than S45A, caused a more severe phenotype that better mimicked the *per*^*SLIH*^ phenotype resulting from a S45Y substitution (Figure 1B, Table 1 (31)). This could indicate an important structural role for this amino acid, perhaps controlling specific protein-protein interactions not implicated in PER degradation.

Interestingly, our work, combined with previous work in mammals, converge to support the model that CKIδ/ε-mediated phosphorylation of PER proteins and their proteasomal degradation are critical for temperature compensation of the circadian clock. Surprisingly, however, there appear to be important mechanistic differences. Indeed, in *Drosophila*, mutation of the key Serine residue (S47) in the phosphodegron increases temperature compensation, while in mammals mutation of the homologous S478 residue has the opposite effect (Figure 1B, 2B, Table 1 (12,13)). Intriguingly, in mammalian cell culture assays, phosphodegron mutation (S478A in mammals) causes degradation kinetics to become essentially insensitive to temperature (12), while S47A degradation in flies is mildly overcompensated. However, in mammals, wild-type mPER2 degradation kinetics accelerates when temperature drops, while in flies we observed the opposite effect, perhaps because the phosphodegrons, though conserved, are located in different location in the PER and mPER2 proteins. This explains why mutations that both renders PER degradation less sensitive to temperature have opposite effect on temperature compensation.

Finally, it is important to point out that so far, any mutation or manipulation of phosphorylation of key circadian protein in any system only partially disrupts temperature compensation, or even increases it. It is therefore clear that multiple mechanisms are implicated, and recent results by Giesecke and collaborators (20) in flies for example point to an important role for nuclear export. It is therefore clear that temperature compensation is proving to be a complex, systemic process, as first proposed by Sweeney and Hastings in their seminal work on Dinoflagellates.

## Material and Methods

### Fly stocks

Fly stocks were maintained on standard cornmeal agar at 25^0^C under 12hr:12hr light: dark (LD). The following available strains were used *w*^*1118*^, *yw*, FM7a, *w[1118]; PBac{y[+mDint2]=vas-Cas9}VK00027* (BL51324), *y[1] M{Act5C-Cas9*.*P*.*RFP-}ZH-2A w[1118] DNAlig4[169]* (BL58492). Transgenic flies used in Figure 2 were described previously (33,39).

### CRISPR mutagenesis

Single stranded oligos (ssODNs) were injected into either BL51324 or BL58492 from Bloomington. The injected embryos, upon eclosion, were crossed with FM7a balancer flies. F1 progeny was screened with PCR and sequencing for the presence of the desired modifications.. All the resultant lines were backcrossed into the *w*^*1118*^genetic background. For lines generated from BL58492 (S45S# I, S45S# II, S45D #122, #67, S45A 98, S4445A# 105, S45E, S45F #51, S44D, S47D), we determined whether the Cas9 transgene located on the X chromosome had been removed. S45E, S44D, S47D fly lines unexpectedly retained Cas9 even after multiple attempts to remove it. It seems however highly unlikely that Cas9 could somehow interfere with temperature compensation, particularly since in both S45E and S44D temperature compensation was unaffected (Figure 1B).

### ssODN design

A conveniently located PAM site (TGG) was identified using CRISPR fly design. The guide RNA sequence chosen was – 5’AGCCACTGCTGCCGGAGGAGTGG. The guide RNAs were cloned into plasmid pCFD3 (Addgene, Cat no: 49410). Silent modifications (AGC-AGT) were introduced into the seed sequence to prevent re-cutting and for subsequent molecular screening.

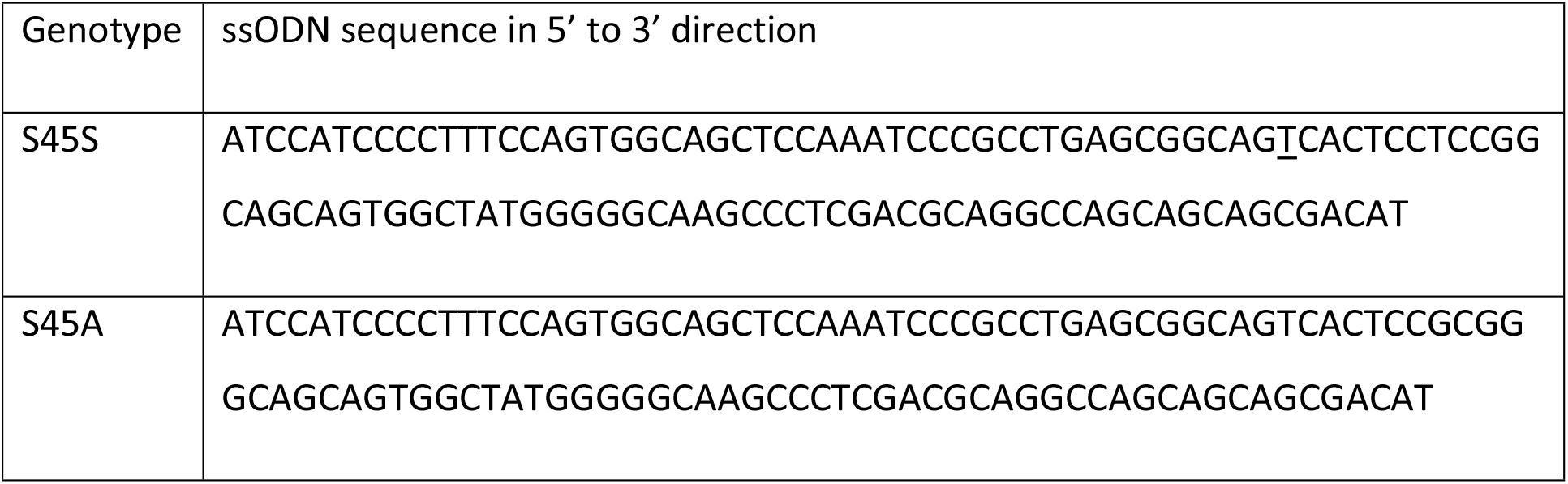
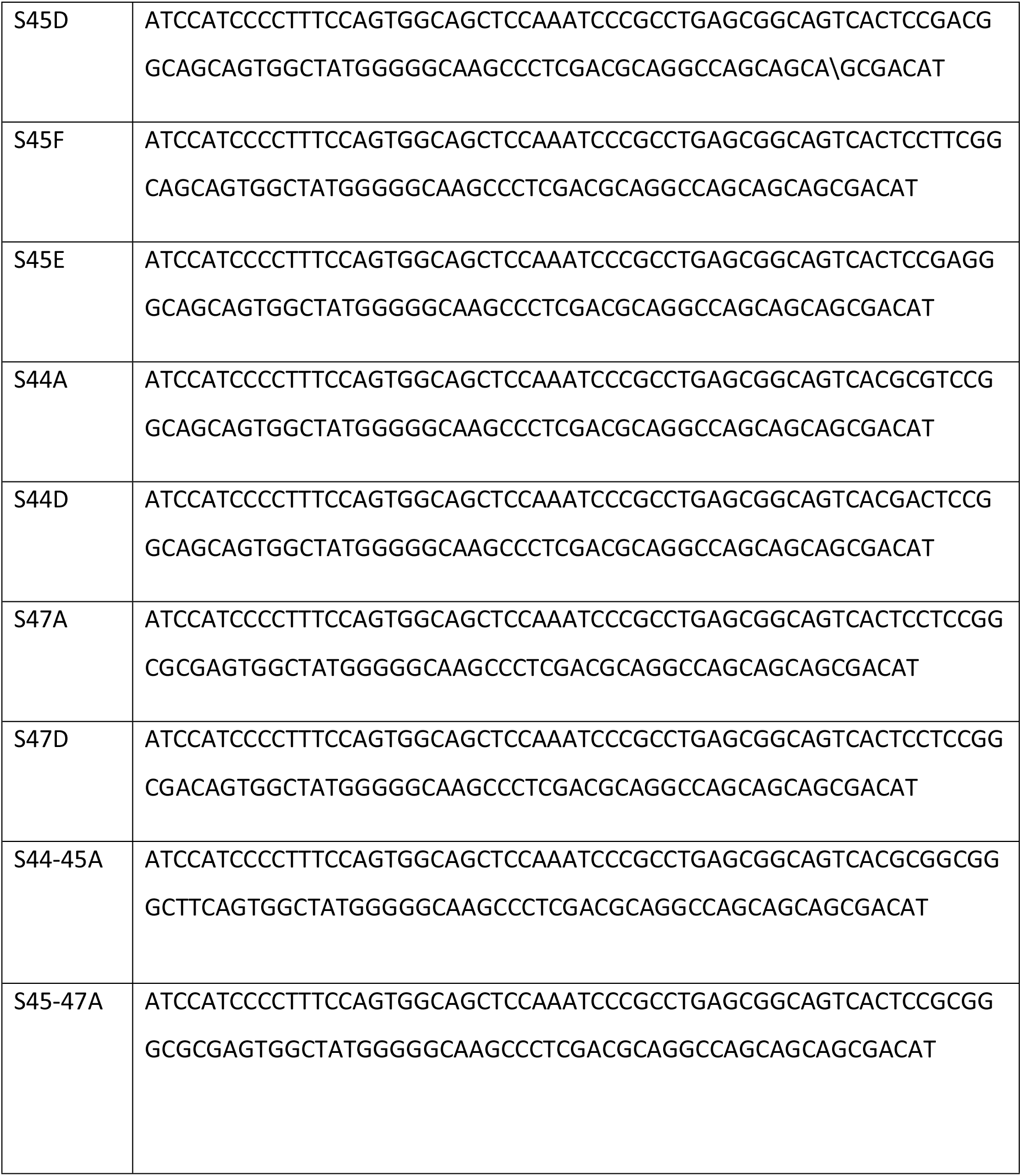

**Primers used**

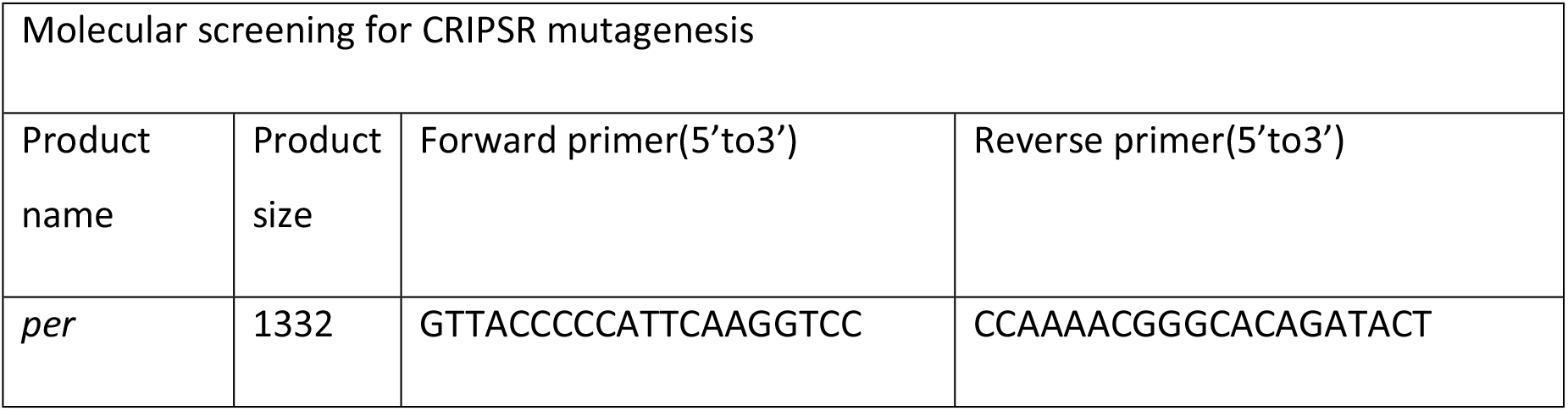
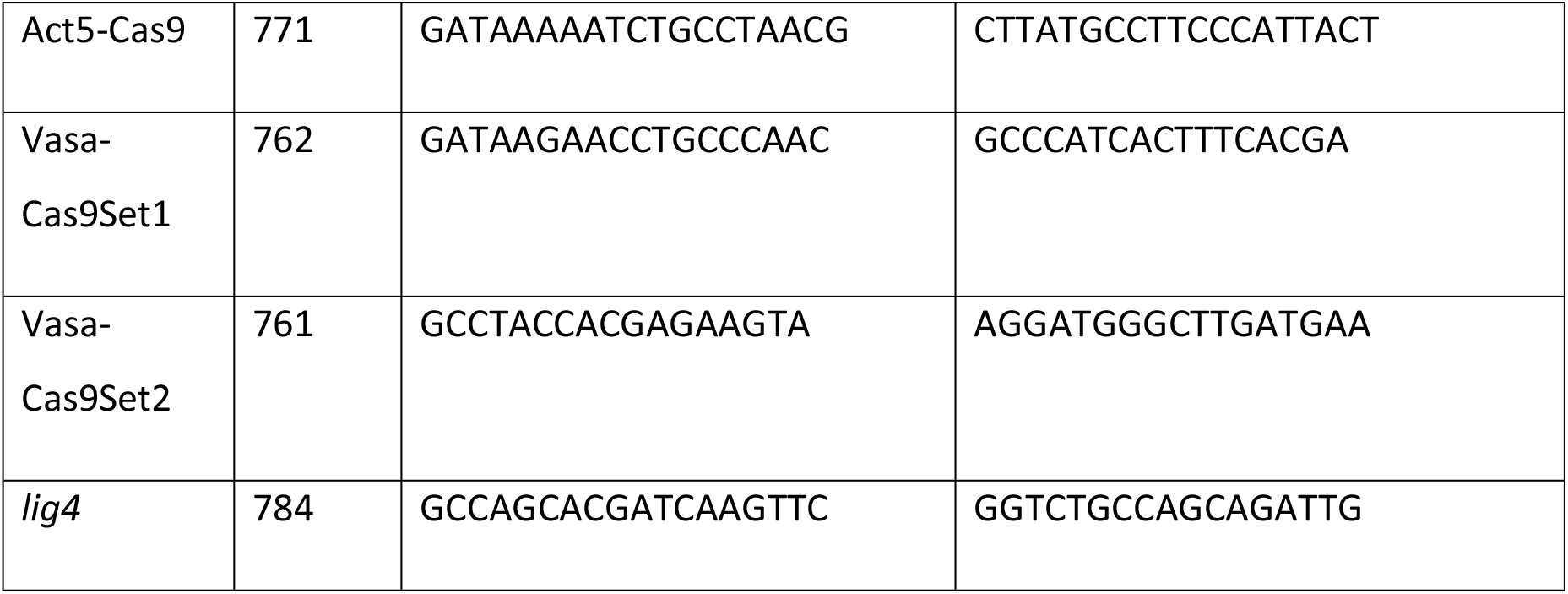

### Circadian locomotor activity monitoring and analysis

2-5 days old male flies were loaded into behavior tubes. Flies were entrained at the defined temperature for at least 3 days under 12:12 LD cycle and released into constant darkness (DD). Activity was recorded using the DAM (*Drosophila* Activity Monitoring) system (Trikinetics, Waltham, MA, USA) in 136-LL incubators (Percival). Behavior was analyzed and plotted using FaasX software (courtesy of F. Rouyer, Centre National de la Recherche Scientifique, Gif-sur-Yvette, France). Rhythmicity was defined by following the criteria: power >20, width >1.5, using the c2 periodogram analysis. 5 days of DD data were used to determine period. Period of all individual flies under various conditions can be found in supplemental file Figure 1-2 - datasets.xls

### Statistical analysis

Whether difference in period at 29^0^C and 18^0^C were statistically significant was determined by using a two-way ANOVA followed by Sidak’s multiple comparison tests. Since in figure 1 control flies showed a significant period shortening at warm temperatures compared to cold, we did factor this drift prior to run statistical test by subtracting the average period lengthening observed in S45SI, and S45SII (1.03hr) from the period observed at 18^0^C for each genotypes.

For graphical representation in figure 1 and 2, mean difference between 29^0^C and 18^0^C were plotted on the Y axis and Standard Error (SE) of difference were used for error bars. Mean difference and SE of difference were obtained from the Two-way Anova statistics details is Prism. Western blots were analysed using Two-way ANOVA with Tukey’s multiple comparison test.

### *Drosophila* S2 cell culture and transfection

*Drosophila* S2 cells were maintained at 22°C in Schneider’s *Drosophila* medium (Life Technologies (Carlsbad, CA)) supplemented with 10% Fetal Bovine Serum (FBS) (VWR, Radnor, PA). Plasmids expressing pAc-*per*(WT)-V5, pAc-*per*(S45A)-V5, pAc-*per*(S47A)-V5 and pMT-*dbt*-V5 were described in Ko et al. (2002) and Chiu et al. (2008). For all cell culture experiments, S2 cells were seeded at 1 × 10^6^ cells/ml in a 6-well plate and transfected using Effectene (Qiagen, Germantown, MD). For mobility shift assay (Figure 4), S2 cells were co-transfected with 0.2μg of pMT-*dbt*-V5 and 0.8μg of pAc-*per*(X)-V5, where X is either WT, S45A or S47A. *dbt* expression were induced with 500μM CuSO_4_ 36 hours after transfection and cells were moved into incubators with indicated temperature and cells were harvested at 0, 6, 12, 24 hours after induction. Proteins were analyzed by Western Blotting. For cycloheximide (CHX) chase experiment (Figure 5), S2 cells were co-transfected with 0.2μg of pMT-*dbt*-V5 and 0.8μg pAc-*per*(X)-V5, where X is either WT or S47A. *dbt* expression were induced with 500μM CuSO_4_ at 36 hours after transfection. 6 hours after DBT induction, CHX (Sigma) was added at a final concentration of 10μg/ml. Cells were harvested and lysed with EB2 (20mM HEPES pH 7.5, 100mM KCl, 5% glycerol, 5mM EDTA, 1mM DTT, 0.1% Triton X-100, 10μg/ml Aprotinin, 5μg/ml Leupeptin, 1μg/ml Pepstatin A, 0.5mM PMSF, 25mM NaF) at the indicated times. Proteins were analyzed by Western Blotting.

### Western blotting and antibodies

Protein extractions from *Drosophila* S2 cells and adult fly heads, western blotting, and image analysis was performed as previously described (Chiu et al., 2008; Cai et al., 2021). 2-5 day old flies were entrained to 12:12 light-dark cycle for 3 days at indicated temperature. On the 4^th^ day of LD, flies were collected at the indicated time points. ∼40 fly heads for each time point were extracted with RBS buffer (20mM HEPES pH7.5, 50mM KCl, 10% glycerol, 2mM EDTA, 1mM DTT, 1% Triton X-100, 0.4% NP-40, 10μg/ml Aprotinin, 5μg/ml Leupeptin, 1μg/ml Pepstatin, 0.5mM PMSF, 25mM NaF). Protein concentration was measured using Pierce Coomassie Plus Assay Reagents (Thermo Fisher Scientific). 2X SDS sample buffer was added and the mixture boiled at 95°C for 5 minutes. Equal amounts of proteins were resolved by polyacrylamide-SDS gel electrophoresis (PAGE) and transferred to nitrocellulose membrane (Bio-Rad, Hercules, CA) using Semi-Dry Transfer Cell (Bio-Rad). Membranes were stained with Ponceau S solution (0.1% Ponceau S, 5% acetic acid) to stain total proteins as loading control. Membranes were then incubated in 5% Blocking Buffer (Bio-Rad) for 40 minutes, incubated with primary antibodies for 16-20 hours. Blots were then washed with 1X TBST for 1 hour, incubated with secondary antibodies for 1 hour, washed again prior to treatment of chemiluminescence Clarity ECL reagent (Bio-Rad). The following percentage of polyacrylamide-SDS gel were used: 6% for PER; 10% for DBT and HSP70.

#### Primary antibodies

α-V5 (Thermo Fisher Scientific) at 1:3000 for PER-V5 and DBT-V5, α-PER (GP5620; RRID:AB_2747405) at 1:2000 for PER, α-HSP70 at 1:10000 for HSP70. Secondary antibodies conjugated with HRP were added as follows: α-mouse IgG (Sigma) at 1:2000 for α-V5 detection, 1:10000 for α-HSP70 detection, α-guinea pig IgG (Sigma) at 1:1000 for α-PER detection. All quantifications are included in supplemental file Figure 3-5-datasets.xls

## Acknowledgments

We thank VInh Phan for technical support, and members of the Emery, Weaver and Anaclet labs for helpful discussions. This work was supported by MIRA award 1R35GM118087 from the National Institute of General Medicine Sciences (NIGMS) to PE, and R01 DK124068 from the National Institute of Diabetes and Digestive and Kidney Diseases (NIDDK) to JCC.

